# Phytoremediation of heavy metals in spent engine oil-polluted soil by *Senna alata* L.

**DOI:** 10.1101/532887

**Authors:** C. Ugwu Emmanuel, O. Nwadinigwe Alfreda, C. Agbo Benita

**Affiliations:** Plant Science and Biotechnology, University of Nigeria, Nsukka, Nigeria.

**Keywords:** Phytoremediation, *Senna alata*, spent engine oil, heavy metals

## Abstract

*Senna alata* L. was used to remediate heavy metals in soil polluted with spent engine oil (SEO). One hundred and twenty polythene bags filled with 20 kg of soil each were separated into parts A and B. Part A contained *S. alata* seedlings while part B had no plant. They were set up in completely randomized design. To simulate spillage, 0.15% v/w, 0.75% v/w and 3.75% v/w concentrations of SEO were used to pollute soil planted with seeds of *S. alata*, 57 days after planting. These treatments were repeated in soil without seeds. Control had no pollution. Heavy metal analyses of SEO, vegetated and non-vegetated soil, leaves, stems and roots of *S. alata* were determined using Flame Atomic Absorption Spectroscopy, 106 days after pollution. Vegetative and reproductive parameters were also determined. Copper, Lead, Zinc, Iron and Aluminium were detected in SEO. Concentrations of heavy metals in vegetated soils were significantly (P < 0.05) less than those of non-vegetated soils. Cu accumulation in stem was significantly (P < 0.05) higher than those of leaves and roots. Zn and Al were significantly (P < 0.05) higher in root than those in leaves and stems. Fe and Pb were significantly (P < 0.05) higher in leaves than in stems and roots. Aerial roots were formed by *S. alata* which increased significantly with increase in concentrations of SEO applied. However, many vegetative parameters such as plant height, number of pinnules, number of roots, leaf area and stem circumference increased significantly (P < 0.05) but some reproductive parameters such as number of inflorescence and dry weight of seeds decreased after pollution. Hence, *S. alata* is suitable for phytoremediation and in particular, phytoaccumulation of heavy metals in SEO contaminated soil.

## Introduction

Increasing number of machines and automobile importation and usage, especially in developing countries has resulted in excessive use of various refined crude oil products such as engine oil, diesel, gasoline etc. Unfortunately, many who utilize these hydrocarbon containing substances have little or no knowledge of the adverse effect of indiscriminate disposal of the used products on the ecosystem. Spent engine oil (SEO) is usually obtained after servicing and subsequent draining from automobile and generator engines by motor mechanics and much of this oil is poured into the soil, gutters, water drains, open plots and farms(1). This indiscriminate disposal of SEO adversely affects plants, microbes and aquatic lives(2). Reference (3) reported that the ratio of the microbial biomass carbon to oxidizable carbon content and basal respiration dropped down significantly on heavily heavy metal contaminated sites. Heavy metals may be retained in the polluted soil from season to season(4), and may lead to build up of essential and non-essential elements in soil which eventually translocated into plant tissues(5). Although heavy metals in low concentrations are essential micronutrients for plants(6), but at high concentrations, they may cause metabolic disorder and growth inhibition for most plant species(7). One of the most deleterious effects induced by heavy metal exposure in plants is lipid peroxidation, which can directly cause biomembrane deterioration(7). Reference (8) reported a significant negative reduction in growth and development and yield of sunflower grown in soil contaminated with heavy metals. Reference (9) also reported a significant decrease in roots and shoot lengths, leaf number, dry and fresh weight of root, shoot and leaf of *Spinacia oleracea* and *Amaranthus caudatus* in heavy metal polluted soil. Also, reference (10) reported a significant decrease in root and shoot lengths of Indian mustard at higher concentrations of heavy metals in a contaminated soil. However, plants and associated microbes have been found to be effective in remediation of heavy metal polluted site(11). Some plants have developed a very potential mechanism to combat such adverse environmental heavy metal toxicity problems(7). These plants might have produced low molecular weight thiols that show high affinity for toxic metals. The most important/critical low molecular weight biological thiols are glutathione (GSH) and cysteine(7). GSH usually elevates serine acetyltransferase activity, which is crucial for detoxification of heavy metals(12). Phytochelatins (PCs) are another small cysteine-rich polypeptide that form complexes with toxic metal ions in the cytosol and subsequently transport them into the vacuole(13), hence, protecting plant tissues from the deleterious effect of heavy metals. However, the ability to accumulate heavy metals varies significantly between species and among cultivars within species, since different mechanisms of ion uptake, metal accumulation, exclusion and compartmentalization are operative in various plant species, based on their genetic, morphological, physiological and anatomical characteristics(14). Reference (15) reported the existence of differential regulation mechanisms on the plasma membrane of *Thlaspi caerulescens* that resulted in the storing of heavy metals in the root vacuoles and thus became unavailable for loading into the xylem and subsequent translocation to shoots. Hence, in phytoremediation, heavy metals are removed by plants in a cost-effective and environmentally friendly manner with reduced exposure to human, terrestrial and aquatic lives.

*Senna alata* (L.) Roxb. (syn. *Cassia alata* L.), commonly known as candle stick senna, wild senna, ringworm cassia and king of the forest, is a medium-sized, ornamental flowering shrub belonging to the Family Fabaceae(16). It is native to Amazon rain forest but spread widely in warm areas of the world especially in the tropical and subtropical regions. So far, no work has been carried out on the phytoremediation of heavy metals in spent engine oil-polluted soil using *Senna alata*. Hence, this work was carried out to bridge this gap.

The objectives of the study were to ascertain the type and quantity of heavy metals that can be removed or accumulated by *S. alata* in soil contaminated with spent engine oil and to determine the vegetative and reproductive parameters of *S. alata* growing on different concentrations of spent engine oil.

## Materials and methods

Black perforated soil bags were filled, each with 20 kg of top soil collected at a depth of 10 cm from Botanic Garden, University of Nigeria, Nsukka. Top soil was mixed with poultry manure at a ratio of 6:2 before bagging. Each of 60 soil bags was planted with a seed of *Senna alata* (vegetated soil), while the other 60 bags were non-vegetated. To simulate spillage, 15 soil bags were polluted with 30 ml (0.15% v/w) of spent engine oil (SEO), 57 days after planting (DAPL). Pollution was repeated with 150 ml (0.75% v/w) and 750 ml (3.75% v/w) of spent engine oil separately, instead of 30 ml. Both vegetated and non-vegetated bags were polluted in the same manner on the same day. Control had no spent engine oil pollution. The experiment was carried out in three replicates and arranged in a completely randomized design. The bags were displayed under the sunlight and watered by rainfall.

### Heavy metal analysis

A sample of SEO, all polluted and unpolluted vegetated and non-vegetated soil samples, leaves, stems and roots of *S. alata* were analyzed for accumulation of heavy metals using flame atomic absorption spectroscopy (FAAS), 106 days after pollution (DAP). Samples were dried at 45°C using Memmert 854 Schwabach oven and crushed into fine powder. One gram was heated for 8 hours in a furnace and cooled in a desiccator. Five ml of trioxonitrate (v) acid (HNO_3_) solution was added to the ash, evaporated to dryness on a hot plate, returned to the furnace and heated at 400°C for 15-20 minutes until perfect greyish-white ash was obtained and allowed to cool in a desiccator. Fifteen ml of hydrochloric acid (HCl) was added to the ash to dissolve it. The solution was filtered into 100 cm^3^ volumetric flask and made up to 100 cm^3^ with distilled water(17). SEO sample was also prepared by digestion method. This was done by putting 2 g into a digestion flask, followed by addition of 20 ml of acid mixture (650 ml conc. HNO_3_, 80 ml perchloric acid and 20 ml sulfuric acid) and heating until a clear solution was obtained. Hexane was added to the flask up to the mark of 25 ml(18). Flame atomic absorption spectroscopic (FAAS) analysis was carried out according to the method adopted by(17). A series of standard metal solutions in the optimum concentration range were prepared by diluting the single stock element solution with water containing 1.5 ml conc. HNO_3_/liter. A calibration blank was also prepared using all the reagents except for the metal stock solution. The sample was aspirated into the flame using Varian AA240 Atomic Absorption Spectrophotometer (AAS) and atomized when the AAS’s light beam is directed through the flame into the monochromator. The atomized sample was directed onto a detector that measured the amount of light absorbed by the atomized element in the flame. Calibration curve for each metal was prepared by plotting the absorbance of standard versus their concentration using Spectra AA scan (PC/Window 7) software.

### Determination of vegetative and reproductive parameters of *Senna alata*

The vegetative parameters of the plant namely, plant height, number of leaves per plant, number of pinnules per leaf, leaf area according to (19) and stem circumference were determined before pollution, that is, at 56 DAPL. After pollution, the same vegetative parameters in addition to the number of roots, root length and root circumference were measured at 163 DAPL. As regards the reproductive parameters, number of inflorescences per plant, flowers, pods, seeds and dry weight of seeds were determined at maturity (at 294 DAPL).

### Data analysis

Data collected from FAAS, vegetative and reproductive parameters were analyzed using (20) to generate the means and variance at P ≤ 0.05. Mean of each parameter was compared using Fisher’s Least Significance Difference. T-test was used to compare vegetative parameters before and after pollution.

## Results and discussion

### Heavy metals

Heavy metals namely, copper (Cu), lead (Pb), iron (Fe), zinc (Zn) and aluminium (Al) were detected in SEO and in all polluted vegetated and non-vegetated soil samples (Table 1) and they generally increased with increase in concentrations of SEO applied. Concentration of each heavy metal in SEO was significantly (P < 0.05) higher when compared with those in polluted vegetated and non-vegetated soils. Also, heavy metal concentrations in polluted non-vegetated soils were significantly (P < 0.05) higher than those in polluted vegetated soils for all treatments.

**Table 1:**
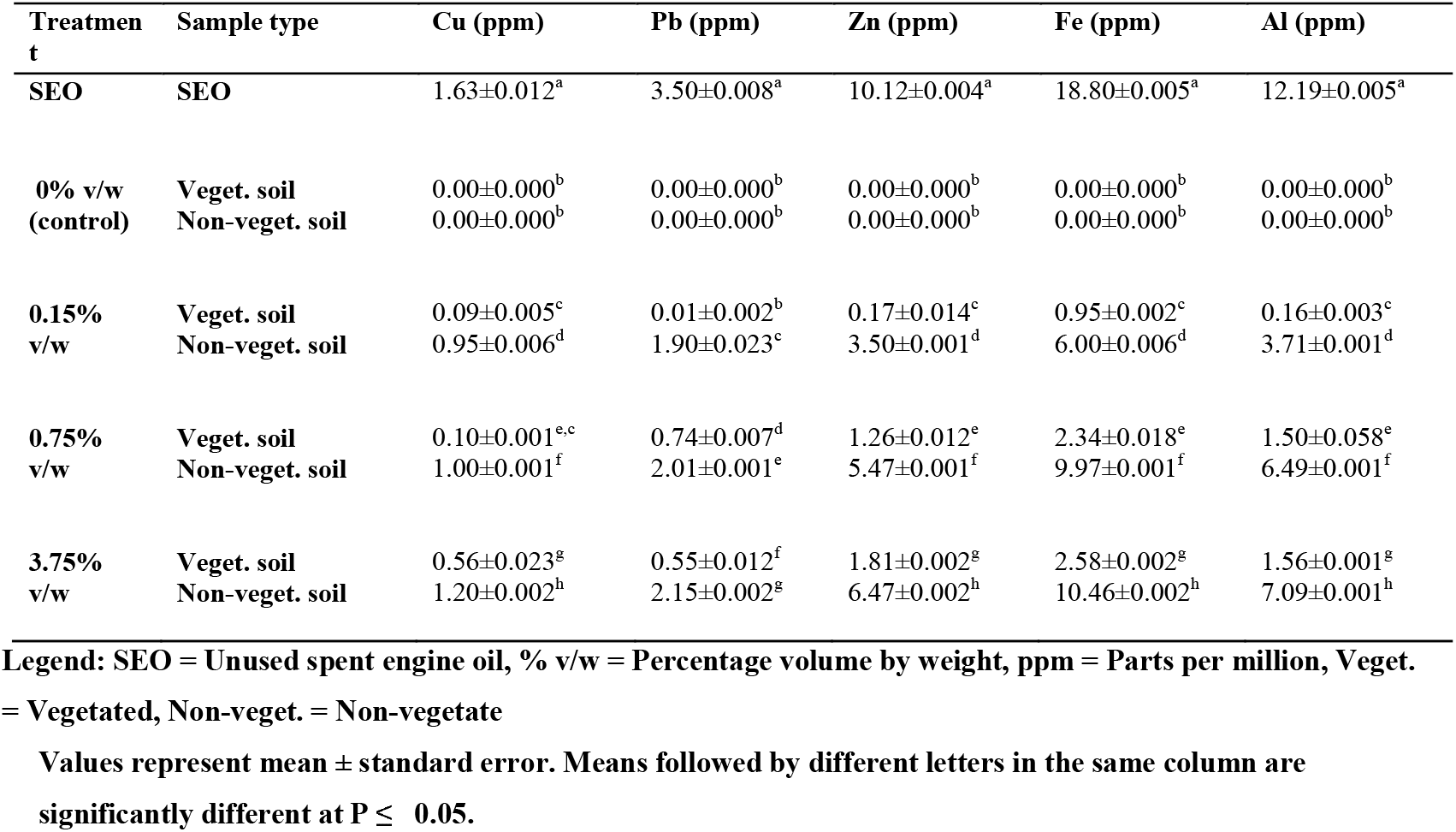
Composition and quantity (ppm) of heavy metals in vegetated and non-vegetated soils of *Senna alata* polluted with spent engine oil.

| Treatment | Sample type | Cu (ppm) | Pb (ppm) | Zn (ppm) | Fe (ppm) | Al (ppm) |
| --- | --- | --- | --- | --- | --- | --- |
| SEO | SEO | 1.63±0.012<sup>a</sup> | 3.50±0.008<sup>a</sup> | 10.12±0.004<sup>a</sup> | 18.80±0.005<sup>a</sup> | 12.19±0.005<sup>a</sup> |
| 0% v/w<br>(control) | Veget. soil | 0.00±0.000<sup>b</sup> | 0.00±0.000<sup>b</sup> | 0.00±0.000<sup>b</sup> | 0.00±0.000<sup>b</sup> | 0.00±0.000<sup>b</sup> |
|  | Non-veget. soil | 0.00±0.000<sup>b</sup> | 0.00±0.000<sup>b</sup> | 0.00±0.000<sup>b</sup> | 0.00±0.000<sup>b</sup> | 0.00±0.000<sup>b</sup> |
| 0.15%<br>v/w | Veget. soil | 0.09±0.005<sup>c</sup> | 0.01±0.002<sup>b</sup> | 0.17±0.014<sup>c</sup> | 0.95±0.002<sup>c</sup> | 0.16±0.003<sup>c</sup> |
|  | Non-veget. soil | 0.95±0.006<sup>d</sup> | 1.90±0.023<sup>c</sup> | 3.50±0.001<sup>d</sup> | 6.00±0.006<sup>d</sup> | 3.71±0.001<sup>d</sup> |
| 0.75%<br>v/w | Veget. soil | 0.10±0.001<sup>e,c</sup> | 0.74±0.007<sup>d</sup> | 1.26±0.012<sup>e</sup> | 2.34±0.018<sup>e</sup> | 1.50±0.058<sup>e</sup> |
|  | Non-veget. soil | 1.00±0.001<sup>f</sup> | 2.01±0.001<sup>e</sup> | 5.47±0.001<sup>f</sup> | 9.97±0.001<sup>f</sup> | 6.49±0.001<sup>f</sup> |
| 3.75%<br>v/w | Veget. soil | 0.56±0.023<sup>g</sup> | 0.55±0.012<sup>f</sup> | 1.81±0.002<sup>g</sup> | 2.58±0.002<sup>g</sup> | 1.56±0.001<sup>g</sup> |
|  | Non-veget. soil | 1.20±0.002<sup>h</sup> | 2.15±0.002<sup>g</sup> | 6.47±0.002<sup>h</sup> | 10.46±0.002<sup>h</sup> | 7.09±0.001<sup>h</sup> |

**Legend: SEO = Unused spent engine oil, % v/w = Percentage volume by weight, ppm = Parts per million, Veget.**

Appreciable quantities of heavy metals were detected in vegetative parts of *Senna alata* (Table 2).

**Table 2:** Composition and quantity (ppm) of heavy metals that accumulated in root, stem and leaf samples of *Senna alata* polluted with spent engine oil.

| Treatment | Sample type | Cu (ppm) | Pb (ppm) | Zn (ppm) | Fe (ppm) | Al (ppm) |
| --- | --- | --- | --- | --- | --- | --- |
| SEO | SEO | 1.63±0.012<sup>a</sup> | 3.50±0.008<sup>a</sup> | 10.12±0.004<sup>a</sup> | 18.80±0.005<sup>a</sup> | 12.19±0.005<sup>a</sup> |
| 0% v/w<br>(control) | Root | 0.00±0.000<sup>b</sup> | 0.00±0.000<sup>b</sup> | 0.00±0.001<sup>b</sup> | 0.00±0.000<sup>b</sup> | 0.00±0.000<sup>b</sup> |
|  | Stem | 0.00±0.000<sup>b</sup> | 0.00±0.000<sup>b</sup> | 0.00±0.000<sup>b</sup> | 0.00±0.000<sup>b</sup> | 0.00±0.000<sup>b</sup> |
|  | Leaf | 0.00±0.000<sup>b</sup> | 0.00±0.000<sup>b</sup> | 0.00±0.000<sup>b</sup> | 0.00±0.000<sup>b</sup> | 0.00±0.000<sup>b</sup> |
| 0.15% v/w | Root | 0.01±0.000<sup>b</sup> | 0.10±0.001<sup>c</sup> | 1.00±0.002<sup>c</sup> | 0.60±0.001<sup>c</sup> | 1.32±0.012<sup>c</sup> |
|  | Stem | 0.18±0.002<sup>c</sup> | 0.01±0.006<sup>d</sup> | 0.02±0.002<sup>d</sup> | 0.64±0.002<sup>c</sup> | 0.20±0.002<sup>d</sup> |
|  | Leaf | 0.003±0.001<sup>c,b</sup> | 0.15±0.002<sup>e</sup> | 0.05±0.001<sup>e</sup> | 3.67±0.012<sup>d</sup> | 0.19±0.017<sup>e</sup> |
| 0.75% v/w | Root | 0.29±0.012<sup>d</sup> | 0.48±0.001<sup>f</sup> | 2.62±0.002<sup>f</sup> | 0.60±0.001<sup>e,c</sup> | 2.02±0.002<sup>f</sup> |
|  | Stem | 0.56±0.002<sup>e</sup> | 0.11±0.006<sup>g,c</sup> | 0.15±0.001<sup>g</sup> | 1.70±0.173<sup>f</sup> | 0.19±0.003<sup>g</sup> |
|  | Leaf | 0.03±0.023<sup>f</sup> | 1.06±0.002<sup>h</sup> | 0.43±0.001<sup>h</sup> | 4.96±0.003<sup>g</sup> | 0.14±0.001<sup>h</sup> |
| 3.75% v/w | Root | 0.37±0.012<sup>g</sup> | 0.41±0.002<sup>i</sup> | 3.41±0.002<sup>i</sup> | 0.78±0.001<sup>h</sup> | 3.55±0.002<sup>i</sup> |
|  | Stem | 0.68±0.001<sup>h</sup> | 0.24±0.023<sup>j</sup> | 0.28±0.002<sup>j</sup> | 1.80±0.005<sup>i</sup> | 0.25±0.029<sup>j</sup> |
|  | Leaf | 0.08±0.003<sup>i</sup> | 1.66±0.006<sup>k</sup> | 0.58±0.012<sup>k</sup> | 6.74±0.003<sup>j</sup> | 0.20±0.001<sup>k</sup> |

**Legend: SEO = Unused spent engine oil, % v/w = Percentage volume by weight, ppm = Parts per million** **Values represent mean ± standard error. Means followed by different letters in the same column are** **significantly different at P ≤ 0.05.**

The concentration of copper stored in stem was significantly (P < 0.05) higher when compared with those stored in leaves and roots of the plant especially for 0.75 and 3.75 % v/w concentrations. The quantities of copper that accumulated in root and leaf in 0.15% v/w treatment did not vary with control. Zinc and aluminium accumulation in roots were significantly (P < 0.05) higher than those in leaves and stems for all concentrations. Also, concentration of iron and lead in leaves were significantly (P < 0.05) higher than those that accumulated in stems and roots for all concentration

### Vegetative and reproductive parameters

Vegetative parameters (Table 3) were not statistically different between the treatments before pollution and also after pollution (separately).

**Table 3:** Vegetative parameters of *Senna alata* before and after pollution (56 and 163 days after planting, respectively)

| Treatm.<br>(% v/w) | Plant height (cm) |  | No. of leaves |  | No. of pinnules |  | Leaf area (cm<sup>2</sup>) |  | stem circum. (cm) |  |
| --- | --- | --- | --- | --- | --- | --- | --- | --- | --- | --- |
|  | Before<br>pollution | After<br>pollution | Before<br>pollution | After<br>pollution | Before<br>pollution | After<br>pollution | Before<br>pollution | After<br>pollution | Before<br>pollution | After<br>pollution |
| **0** | 36.4±2.4<sup>a</sup> | 225.0±8.3<sup>b</sup> | 13.5±0.6<sup>c</sup> | 27.3±2.5<sup>d</sup> | 9.6±0.3<sup>e</sup> | 18.3±0.3<sup>f</sup> | 240.5±21.5<sup>g</sup> | 1012.6±83.4<sup>h</sup> | 3.8±0.1<sup>j</sup> | 9.7±0.4<sup>k</sup> |
| **0.15** | 38.4±1.6<sup>a</sup> | 215.3±9.9<sup>b</sup> | 13.6±0.7<sup>c</sup> | 26.0±3.0<sup>d</sup> | 9.6±0.4<sup>e</sup> | 17.3±1.1<sup>f</sup> | 251.6±23.3<sup>g</sup> | 992.0±59.3<sup>h</sup> | 3.9±0.08<sup>j</sup> | 9.5±0.4<sup>k</sup> |
| **0.75** | 36.2±2.7<sup>a</sup> | 214.20±7.8<sup>b</sup> | 13.5±0.6<sup>c</sup> | 21.7±2.2<sup>d</sup> | 9.3±0.4<sup>e</sup> | 18.2±0.5<sup>f</sup> | 249.5±25.7<sup>g</sup> | 910.9±60.8<sup>h</sup> | 3.9±0.1<sup>j</sup> | 9.0±0.4<sup>k</sup> |
| **3.75** | 37.8±2.7<sup>a</sup> | 203.4±9.3<sup>b</sup> | 14.5±0.7<sup>c</sup> | 18.3±2.2<sup>d,c</sup> | 9.5±1.2<sup>e</sup> | 16.7±1.3<sup>f</sup> | 240.6±27.2<sup>g</sup> | 842.6±92.1<sup>h</sup> | 3.9±0.1<sup>j</sup> | 8.6±0.4<sup>k</sup> |

However, in comparison, these vegetative parameters determined before pollution were significantly (P < 0.05) lower than vegetative parameters measured after pollution within the treatments. The only exception was the number of leaves produced in 3.75% v/w treated plants whose value did not differ before and after pollution.

As regards the root parameters, results showed that 3.75% v/w treatment significantly (P < 0.05) produced the highest number of roots (Table 4).

**Table 4:** Root parameters of *Senna alata* 163 days after planting.

| Treatment (%v/w) | Root number | Root length (cm) | Root circumference (cm) | Initial no. of aerial root (56 DAP) | Final no. of aerial root (106 DAP) |
| --- | --- | --- | --- | --- | --- |
| **0 (control)** | 61.2±6.009<sup>a</sup> | 56.52±2.107<sup>b</sup> | 1.300±0.058<sup>c</sup> | 0.67±0.159<sup>h</sup> | 0.00±0.000<sup>a</sup> |
| **0.15** | 65.2±2.056<sup>a</sup> | 58.23±0.716<sup>b</sup> | 1.150±0.087<sup>c</sup> | 3.13±0.904<sup>h</sup> | 6.40±1.027<sup>b</sup> |
| **0.75** | 60.5±4.406<sup>a</sup> | 55.92±2.673<sup>b</sup> | 1.125±0.025<sup>c,d</sup> | 7.53±0.867<sup>e</sup> | 13.60±1.287<sup>c</sup> |
| **3.75** | 77.5±3.227<sup>b</sup> | 54.67±2.175<sup>b</sup> | 1.050±0.029<sup>d,e</sup> | 27.87±1.973<sup>f</sup> | 55.07±2.306<sup>d</sup> |

For mean root circumference, control and 0.15% v/w were significantly (P < 0.05) thicker than 3.75% v/w treatment. *Senna alata* also produced aerial roots. Initial (at 56 DAP) and final (106 DAP) number of aerial roots increased significantly (P < 0.05) with increase in concentrations of SEO applied. For each treatment, the final number of aerial roots was significantly (P < 0.05) higher than the initial number.

There were no differences between control and polluted plants as regards the number of flowers, pods and seeds produced by *S. alata* (Table 5).

**Table 5:** Reproductive parameters of *Senna alata* 294 days after planting.

| Treatm.<br>(% v/w) | No. of inflors. | No. of flowers | No. of pods | No. of seeds | Dry wt. of seeds<br>(g) |
| --- | --- | --- | --- | --- | --- |
| 0 | 7.8±0.917<sup>a</sup> | 155.1±14.02<sup>b</sup> | 51.4±4.672<sup>c</sup> | 30.7±1.075<sup>d</sup> | 49.0±4.392<sup>e</sup> |
| 0.15 | 8.5±0.898<sup>a</sup> | 144.3±14.20<sup>b</sup> | 43.8±5.337<sup>c</sup> | 24.7±1.350<sup>d</sup> | 34.6±5.418<sup>e,f</sup> |
| 0.75 | 5.1±0.900<sup>b,c</sup> | 131.5±23.76<sup>b</sup> | 39.7±9.175<sup>c</sup> | 21.4±4.951<sup>d</sup> | 37.5±10.47<sup>e,f</sup> |
| 3.75 | 5.2±1.104<sup>a,c</sup> | 97.4±17.06<sup>b</sup> | 27.7±6.112<sup>c</sup> | 24.0±4.266<sup>d</sup> | 21.3±4.764<sup>f</sup> |

**Legend: Treatm. = Treatment; Inflors. = Inflorescences; No. = Number; Wt. = weight**
**Values represent mean ± standard error. Means followed by the same letters in the same column are not significantly different at $P \leq 0.05$**

Mean number of inflorescences for the control was significantly (P < 0.05) more than that of 0.75% v/w but the same with the rest of the treatments. Dry weight of seeds polluted with 3.75% v/w was significantly (P < 0.05) less than that of control but the same with 0.15% v/w and 0.75% v/w treatments.

## Discussion

In the present study, quantities of heavy metals that remained in the soil in each treatment increased with increase in the concentrations of SEO applied. This implies that removal efficiency of heavy metals decreased with increase in concentrations of SEO applied. This agreed with the findings of (21) who reported that phytoremediation efficiency of metals greatly depends on the concentrations of such metals in solutions; the higher the concentration of metals in solution, the lower the removal efficiency. However, the concentrations of heavy metals accumulated in the vegetative parts of the plant increased with increase in concentration of SEO applied. This agreed with the findings of (22) who reported that the amount of heavy metal accumulation is dose-dependent, as the higher the concentration, the higher the quantity accumulated. Moreover, the concentrations of heavy metals in some vegetative parts of *S. alata* were higher than the concentrations in the vegetated soil. This indicates that the plant organs might have taken up and accumulated these heavy metals. This disagreed with the findings of (23) who reported low to zero removal of heavy metals by water hyacinth. Therefore, higher concentrations of these heavy metals in the vegetative parts of *S. alata* showed that some plants can hyper accumulate these substances and consequently, affect other organisms along the food chain through bioaccumulation. However, the quantities of each of the heavy metals in *S. alata* in each treatment were below the standard allowable limits for daily dietary intake and medicinal plants according to toxicology report/world health organization respectively - (Cu – 60 mg/day; 10 ppm, Zn – 15 mg/day; 50 ppm, Pb – 10 ppm, Fe – 15 mg/day; 20 ppm and Al – 60 mg/day)(24)-(25). Even though the concentrations of these heavy metals in some vegetative parts of *S. alata* were higher than those in the polluted vegetated soil, yet the SEO with heavy metals did not have adverse effects on the vegetative growth and development of the plant, since the plants grew and developed well when compared with the control. This shows that the plant can be used to phytoremediate heavy metals in SEO contaminated soil. This is in contrast to the report of (7) that heavy metals in plants lead to biomembrane deterioration due to lipid peroxidation; consequently, leading to poor vegetative growth, death of seeds and plants(1)-(10)-(9). Hence, *S. alata* may perhaps possess substances that might have formed complexes with toxic metals(7), which might have protected the plant from the deleterious effect of heavy metals. Since *S. alata* accumulated Cu, Pb, Zn, Fe and Al in the leaves, stems and roots, the plant can be used as a hyper accumulator for phytoremediation of some heavy metals. In this way, these heavy metals are removed from the environment. In a similar way (26) reported that *Cassia tora* accumulated Fe, Zn, Cu and Pb at high concentrations and it could be used as hyper accumulator plant for bioremediation. Moreover, Cu was higher in the stems than in the other parts of the plant in the present work. Pb, Zn and Fe were found in larger quantities in the leaves and roots than in the stem while Al was found in large quantities in the roots. These suggest that different mechanisms of metal accumulation, exclusion and compartmentalization exist in various plant organs and species(14) (Mohammad *et al*., 2008). Cosio *et al*. (2004) (15)found that more Zn was stored in the root vacuole of *Thlaspi caerulescens* and thus became unavailable for loading into the xylem because of the existence of differential regulation mechanism on the plasma membrane. Gupta and Sinha (2008) (26)reported that *Cassia tora* accumulated high concentrations of Pb, Cu, Fe and Zn in their leaves and roots.

Since most of the vegetative parameters were the same as the control, 106 days after pollution and they were significantly higher than those obtained before pollution, it means that SEO or heavy metal pollution had no adverse effects on the growth of the plant. Rather, they enhanced the growth and productivity of the plant. Hence, the plant can be used for phytoremediation, because for phytoremediation to take place, the plant must survive and grow in the contaminated medium(27). This is in contrast to the findings of (8) who reported a significant negative reduction in growth, development and yield of sunflower grown in soil contaminated with heavy metals. Number, length and circumference of roots for all the treatments were the same with those of the control. Therefore, they were not adversely affected by SEO or heavy metal pollution. This in contrast to the findings of (10) who reported a significant decrease in root length of Indian mustard (*Brassica juncea*) in soil contaminated with heavy metals. Number of aerial roots per plant increased with increase in SEO concentration. This means that heavy metals in SEO did not inhibit root production. Therefore, spent engine oil with heavy metals might have induced physiological or genetic changes that initiated aerial root formation which is not common in this plant. This is in contrast with the findings of (9) who reported a significant decrease in the number of roots of *Spinacia oleracea* and *Amaranthus caudatus* in heavy metal polluted soil. Reference (28) reported a significant reduction in root cross-section of *Vigna radiata* in soil contaminated with petroleum hydrocarbons.

The number of inflorescences and dry weight of seeds decreased significantly (P < 0.05) with increased concentration of SEO applied. However, the number of flowers, pods and seeds of *S. alata* were the same with the control. Reference (29) reported early flowering but significant reduction in number of flowers, pods and yield of *Cajanus cajan* in soil polluted with cadmium. Reference (1) also reported that grain yield of maize in SEO contaminated soil were adversely affected.

## Conclusion

*Senna alata* is a suitable plant for the phytoremediation of soil polluted with SEO. The plant can also be used for the absorption and phytoaccumulation of heavy metals such as Cu, Pb, Zn, Fe and Al from SEO. These metals did not have negative effect on *S. alata* rather they accumulated in the leaves, stems and roots of the plant. Therefore, *S. alata* can be used as a hyper accumulator plant for phytoremediation. In this way, these heavy metals are removed from the environment. Planting of this perennial shrub will decontaminate these pollutants in oil-polluted areas, aerate the environment during photosynthesis and provide aesthetic value to the environment due to its beautiful flowers. Bioaccumulation of heavy metals along food chain should be curbed through recycling of SEO rather than indiscriminate pouring into the environment which might have negative impacts on biodiversity.

## Recommendations

One major discovery in this work was the aerial root formation which is not common in this plant. More work is needed to unravel the causes and consequences of such incident. It might be as result of mutation or mere physiological/anatomical changes induced by heavy metals or hydrocarbons present in spent engine oil or combined interactions of heavy metals and hydrocarbons; or unknown substance(s) in spent engine oil. There is also need to device means of recycling this waste instead of disposing indiscriminately, since unidentified effects of this pollutant exist, which might pose serious risk to the present and future generations.

## Supporting information

Plates showing vegetative parts of *S. alata* before pollution, after pollution and the aerial roots formed 56 days and 106 days after pollution.

## Figure

**Plate 1:**
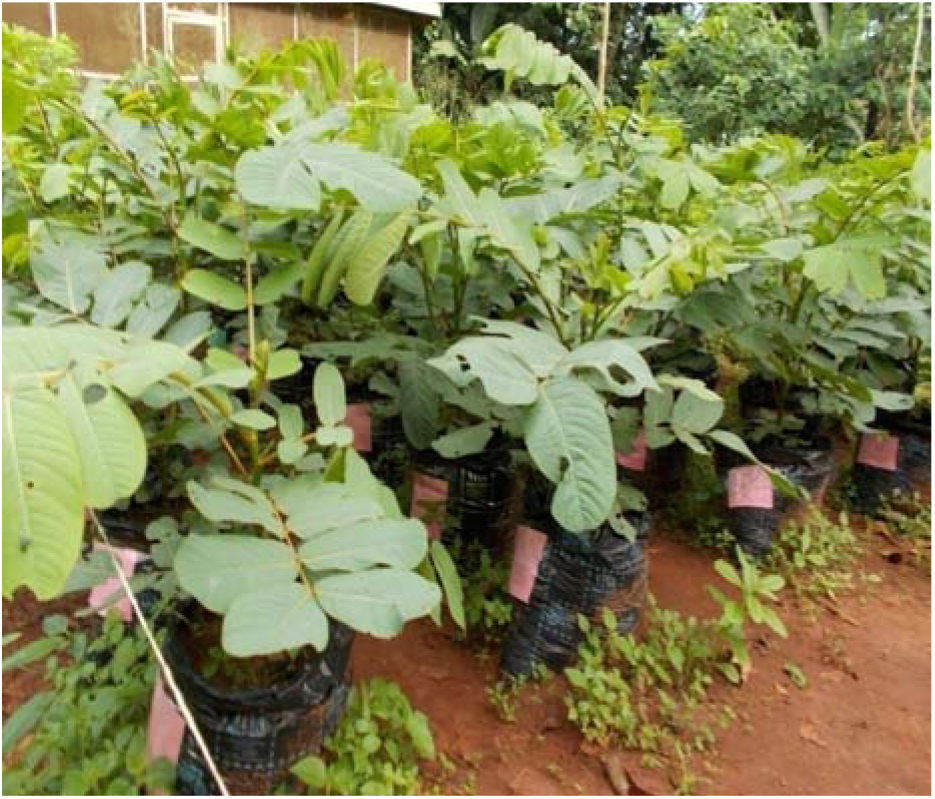
*Senna alata* seedlings, 56 days after planting (DAP)

**Plate 2:**
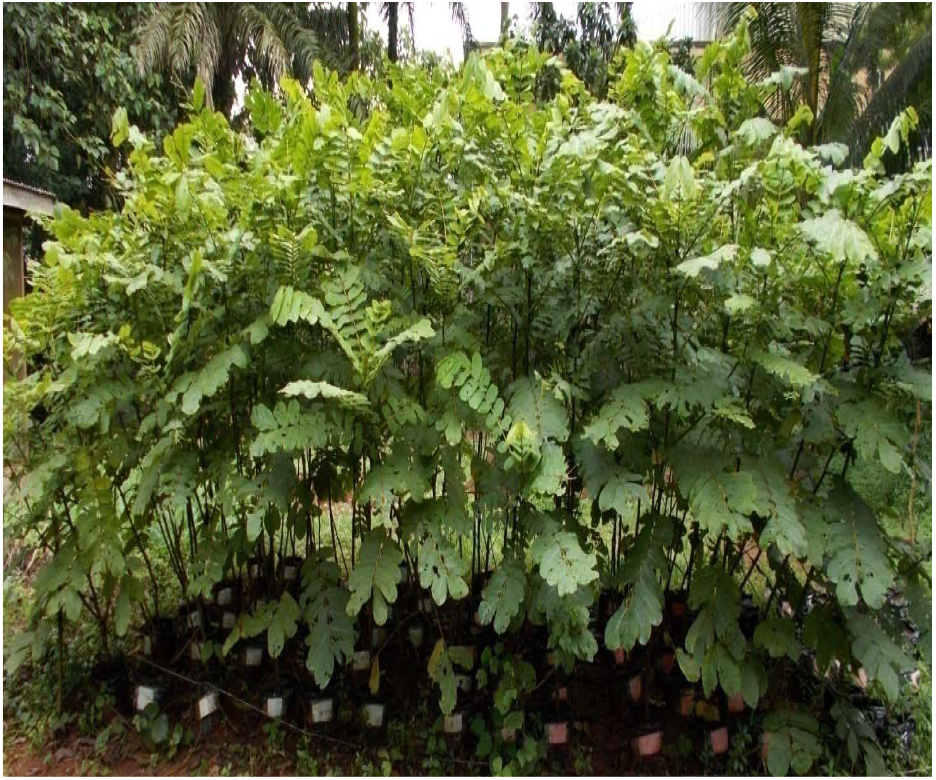
*Senna alata* plants 163 days after planting (106 days after pollution)

**Plate 3:**
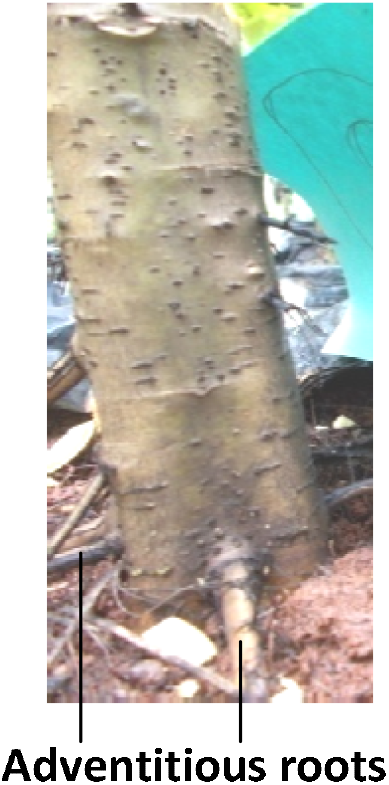
*Senna alata* produced adventitious roots that entered the soil instead of aerial roots (Control)

**Plate 4:**
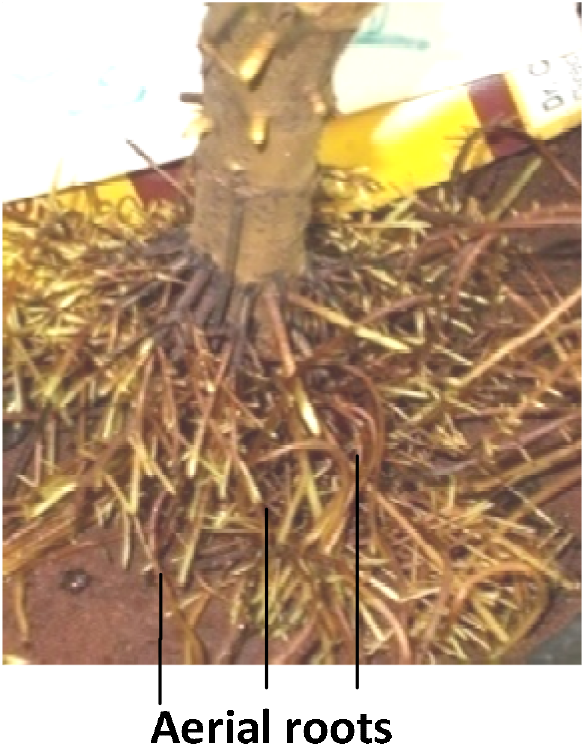
Numerous aerial roots produced by *Senna alata* (56 days after pollution with 3.75% v/w)

**Plate 5:**
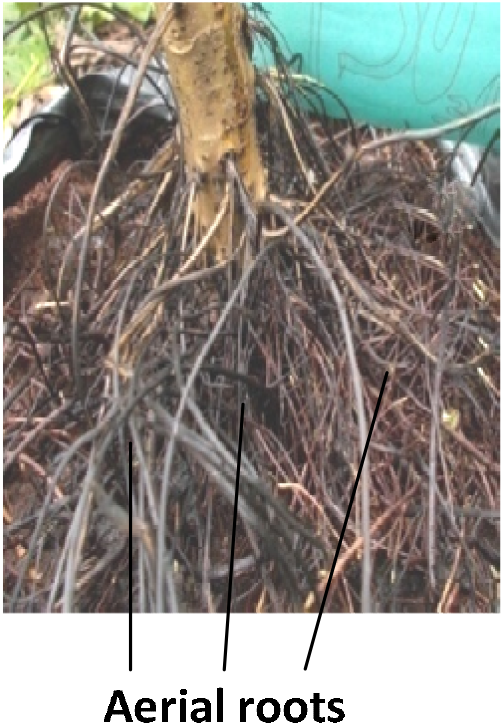
Numerous aerial roots produced by *Senna alata* (106 days after pollution with 3.75% v/w)

